# AFid: A tool for automated identification and exclusion of autofluorescent objects from microscopy images

**DOI:** 10.1101/566315

**Authors:** Heeva Baharlou, Nicolas P Canete, Kirstie M Bertram, Kerrie J Sandgren, Anthony L Cunningham, Andrew N Harman, Ellis Patrick

## Abstract

Autofluorescence is a long-standing problem that has hindered the analysis of images of tissues acquired by fluorescence microscopy. Current approaches to mitigate autofluorescence in tissue are lab-based and involve either chemical treatment of sections or specialized instrumentation and software to ‘unmix’ autofluorescent signals. Importantly, these approaches are pre-emptive and there are currently no methods to deal with autofluorescence in acquired fluorescence microscopy images. To address this, we developed Autofluorescence Identifier (AFid). AFid identifies autofluorescent pixels as discrete objects in multi-channel images post acquisition. These objects can then be tagged for exclusion from downstream analysis. We validated AFid using images of FFPE human colorectal tissue stained for common immune markers. Further, we demonstrate its utility for image analysis where its implementation allows the accurate measurement of HIV-Dendritic Cell interactions in a colorectal explant model of HIV transmission.

**Availability and implementation:** https://ellispatrick.github.io/AFid

## Introduction

Immunofluorescence microscopy (IF) is a powerful tool for simultaneously visualising the localisation of multiple proteins *in situ*. Additionally, several methods have been developed that push the number of parameters visualised in a single section to well beyond traditional 3-4 colour IF^1–7^. This allows for the definition of multiple cell types, complex subsets, and also cellular states *in situ.* Despite these advances the utility of IF, particularly for quantitative measurements, has been hampered by the longstanding issue of autofluorescence.

Autofluorescence is present in all tissues and has many sources including components of structural and connective tissues, cellular cytoplasmic contents and also fixatives used to preserve tissue^8–11^. Autofluorescent substances have their own excitation and emission profiles that can span the entire visible and even infra-red spectrum and therefore significantly overlap with standard microscope excitation/emission filter setups^10^ **(Supplementary Fig. 1**). This presents a major obstacle to image analysis, particularly any kind of automated analyses, as ‘real’ vs ‘autofluorescent’ regions of interest (ROIs) cannot be readily distinguished. An example of this is shown (**Supplementary Fig. 2**) where the accurate quantification of CD3 labelling in human colon tissue is severely hampered by autofluorescent signals.

Several methods have been developed to address the issue of autofluorescence. The oldest and most widely used are chemical methods to quench autofluorescence. These include exposing samples to either UV radiation or a chemical solution prior to or during staining^9,12–14^. Although these methods can be effective, there are several disadvantages including quenching of desired signal from endogenous reporters or fluorescent probes, and also that there is no general recipe with specific protocols required to quench specific types of autofluorescence ^9,14^. However, the primary limitation is that the quenching must take place prior to imaging, so if autofluorescence is detected after image acquisition it is too late to remove it. This can be frustrating as autofluorescence is highly variable between tissue sections.

Digital methods of autofluorescence mitigation have also been developed such as spectral unmixing and algorithmic subtraction of a background reference image acquired either prior to staining^2,15–17^. These methods are robust and have the capacity to resolve signal vs autofluorescence. As such they represent an important pre-processing step to ensure accurate image analysis. However, there are several limitations to these approaches. Spectral unmixing requires the use of specialised instrumentation and proprietary software which limits its generalised use^15^. Similarly background subtraction using a customised filter setup requires a microscope with tuneable filters and expertise beyond that of the average researcher^17^. The alternative background subtraction method requires that entire tissue sections are imaged at a pre-defined resolution prior to staining, whereby the user must perform intensity scaling and pixel-perfect registration, again representing a major hurdle to the average researcher^2^.

Taken together, there is an unmet need for the development of alternative, user-friendly and open-source methods to tackle the longstanding issue of autofluorescence. Based on the long excitation/emission wavelengths of autofluorescence^10^ **(Supplementary Fig. 1**) and the observation that in many cases the majority of interfering autofluorescence is spatially distinct from signal of interest (examples shown in figures throughout this paper), we reasoned that we could develop a post-acquisition approach to identify and exclude autofluorescence thereby improving image analysis accuracy. To this end, we developed ‘Autofluorescence Identifier’ (AFid), an algorithm which is able to detect autofluorescent pixels as discrete objects within multi-channel IF images of tissue. AFid requires only the information from two fluorescent channels, where bright fluorescent ROIs are located and classified as ‘real’ or ‘autofluorescent’ based on measures of pixel correlation, distribution and dynamic range. Identified autofluorescent objects can then be tagged for exclusion from downstream analysis. A key advantage of this method is that it is applied to images post-acquisition and can therefore be used to filter existing image data sets. In this paper we describe the steps of the AFid algorithm, validate its usage on FFPE human colorectal tissue and also demonstrate its utility for image analysis where its implementation allows the accurate measurement of HIV-Dendritic Cell (DC) interactions in a colorectal explant model of HIV transmission.

## Results

### Algorithm overview

The steps for the AFid algorithm are summarised in **Fig. 1.** First, thresholds are applied to the two fluorescent channels and an ‘intersection mask’ is created to detect the ROIs that are fluorescing in both channels (**Fig. 1a, left**). Second, we then measure ROIs in the ‘intersection mask’ for multiple textural features **(Fig. 1a, middle)**. To select these features, we make a key assumption that the fundamental topology of pixel intensities for an autofluorescent ROI is conserved across channels. This makes sense as sources of autofluorescence tend to have long excitation and emission profiles. As such, any measure of pixel behaviour within an ROI will be linearly correlated across channels (**Supplementary Fig. 3a-c**). Therefore, to identify autofluorescence we measure multiple features including pixel correlation (Pearson’s correlation coefficient), dynamic range (Standard Deviation) and distribution (Kurtosis). Third, ROIs can be clustered using the textural features as inputs to identify a distinct cluster with high correlation values that consists mainly of autofluorescent ROIs (**Fig. 1a, right**). Here we have used a *k*-means clustering algorithm with automated choice of *k*. Finally, these autofluorescent ROIs can be excluded from downstream analysis or can be subtracted from the raw images for visualisation (**Fig. 1b,c**).

**Fig. 1:**
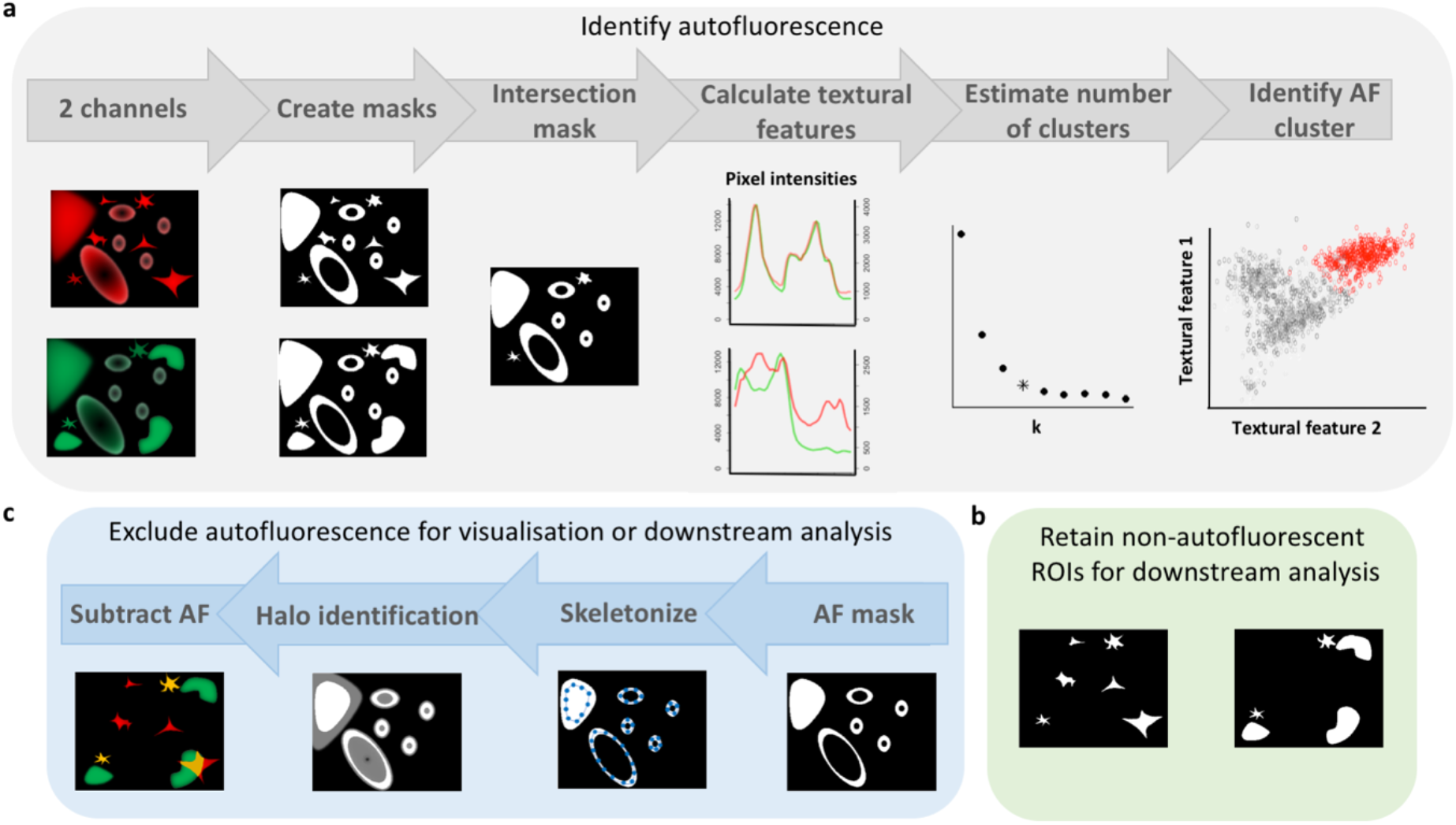
Steps of the AFid algorithm. **a,** *k-means* clustering on a set of textual features of objects in an intersection mask of two channels. Autofluorescent ROIs can then be tagged for exclusion in downstream analysis (**b**), or a custom dilation function can be employed to estimate the perimeter of autofluorescent ROIs, which are then excluded from the image (pixel values set to 0) (**c**).

### Custom dilation function to outline autofluorescent objects

For optimal visualisation, and to aid in downstream analysis, we have also developed a novel algorithm which expands from autofluorescent objects to capture their full perimeter. Due to variations in intensity scale within an image and across different images, conventional thresholding algorithms rarely capture the entire perimeter of autofluorescent ROIs (**Supplementary Fig. 4a-c**). This can represent a limitation for automated autofluorescence exclusion, as several threshold parameters need to be tested beforehand for each image, and then assessed by eye to determine appropriateness. To overcome this limitation we developed a custom dilation function that works in tandem with thresholding to automatically outline the full body of autofluorescent ROIs, regardless of shape and intensity (**Fig. 1c)**. In brief this works by skeletonising ROIs and evenly distributing points throughout the skeleton (**Supplementary Fig. 4d**). We then expand from these points until the gradient of pixel intensities from the ROI boundary outwards begins to increase, indicating the end of the object or the beginning of a neighbouring object (**Supplementary Fig. 4e**). Visually we can see that this method accurately captures the full perimeter of autofluorescent ROIs with minimal effect to neighbouring signals (**Supplementary Fig. 4f**). Further, we quantified the percentage of autofluorescence captured by AFid with and without custom dilation for varying threshold radii and cluster numbers *k* (**Supplementary Figure 4g**). The results show that the custom dilation function identifies a higher percentage of autofluorescent pixels which is stable against variation of threshold radii and cluster number *k*, with minimal impact on neighbouring real signals (with the exception of small *k* values which are typically not selected by AFid). Accordingly, employment of the dilation function allows for efficient autofluorescence detection despite variation in choice of threshold radii. The final result is an image retaining only non-autofluorescent ROIs, which can then be used for visualisation and downstream analysis (**Fig. 1c**).

### Validation of AFid

In order to establish both the efficacy and scope of utility for AFid we tested the algorithm with multiple types of input images. First, we tested whether the markers for detection in each channel could influence autofluorescence identification. To this end we defined three use-cases where input channels contained (1) non-co-expressed markers (CD11c and CD3), (2) a marker expressed on autofluorescent cells (FXIIIA+ Macrophages) and (3) co-expressed markers (CD3 and CD4). These three use-cases are shown for human colorectal tissue where sections were imaged before (**Fig. 2a,d,g**) and after staining (**Fig. 2b,e,h**). The unstained image was used for manual annotation of autofluorescent ROIs providing a ground truth for our classifier, and not used by AFid. We found that in all three use-cases the autofluorescence cluster was highly enriched for autofluorescent ROIs (mean =98.4%, SD = 1.6%, n=9). (**Supplementary Fig. 5a**). We also achieved good coverage identifying on average 96.0% (SD = 4.0%, n=9) of all annotated autofluorescence, with a low false positive rate of 1.7% (SD = 1.7%, n=9) (**Supplementary Fig. 5b**). Pairwise plots for each use-case are shown (**Supplementary Fig. 6–8**), demonstrating g≥ood separation of autofluorescence (yellow) from non-autofluorescence (grey) by *k*-means. The final result of AFid, after applying the custom dilation function is shown (**Fig. 2c,f,i**), demonstrating near complete exclusion of autofluorescence in all three use-cases.

**Fig. 2:**
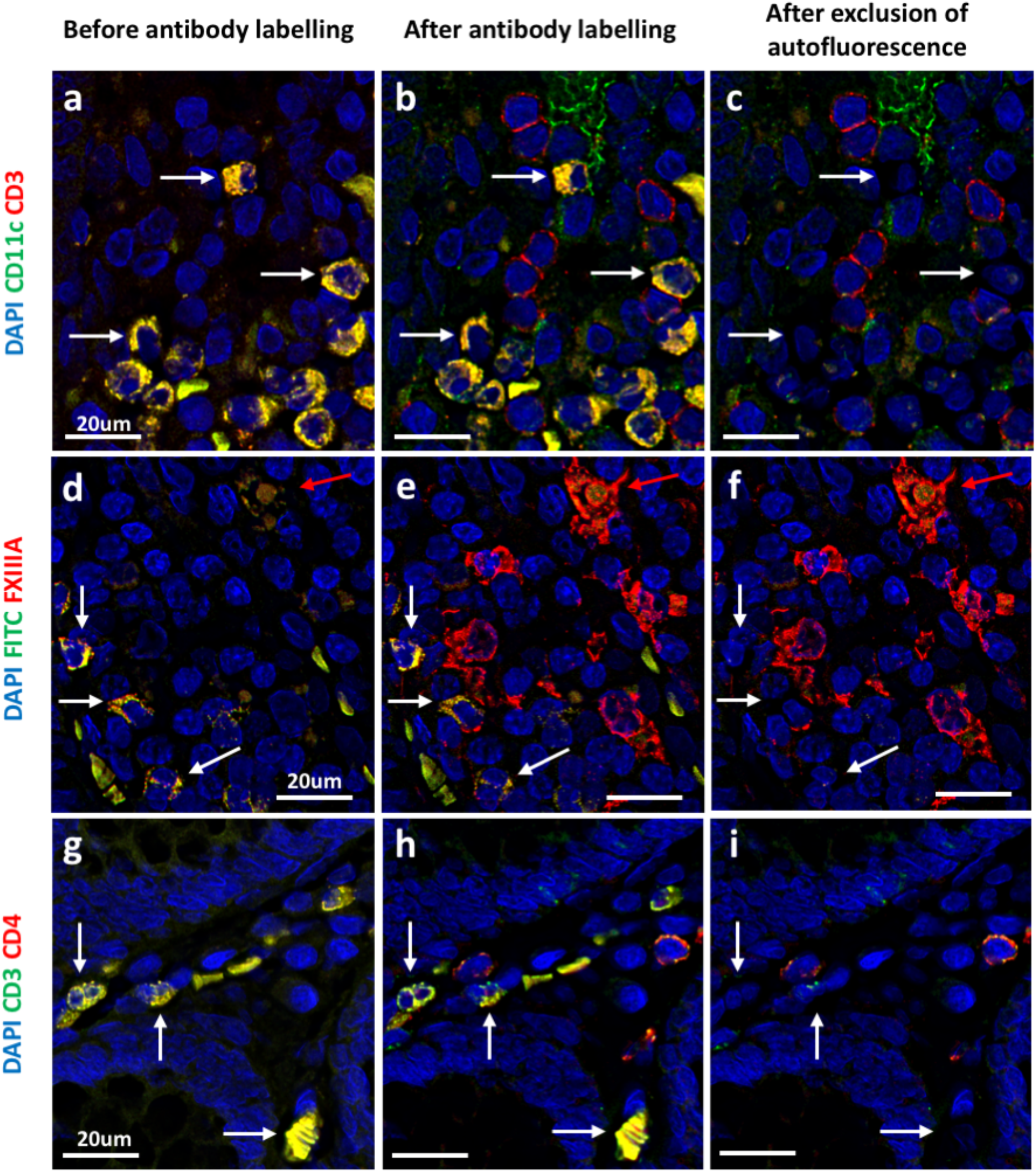
Application of AFid to various staining panels. Sections of fixed human colorectal tissue prior to (**a,d,g**) and after labelling (**b,e,h**) with antibodies targeting the indicated markers. **c,f,i**, labelled images after identification and exclusion of autofluorescencent objects using AFid. White arrows indicate some autofluorescent objects that have been excluded by the algorithm. The red arrow in the middle row indicates an autofluorescent macrophage which was not identified by AFid. In the the middle row, ‘FITC’ is the FITC channel, which was imaged but not used to detect any markers. Images are representative of 3 donors for each staining panel.

We next sought to test AFid on different tissue types and image acquisition conditions. We found that AFid successfully identified autofluorescence across various tissue-types, including heart, skin and brain tissues where autofluorescence is a well-known problem (**Supplementary Fig. 9**). Additionally, AFid could specifically identify autofluorescent objects despite varying image quality, including variations in resolution, (**Supplementary Fig. 10)** and on images both pre- and post- deconvolution (**Supplementary Fig. 11**). Further, our algorithm is compatible with very large images that are inundated with autofluorescence, which allows large data sets with significant noise to be rescued for analysis (**Supplementary Fig. 12**).

### AFid allows accurate assessment of HIV-Dendritic Cell interactions in an explant infection model

Finally, we show that the presence of autofluorescence and its removal can have a major impact on down-stream analysis. In our own studies, we are interested in the early HIV-target cell interactions that occur in human colorectal explants that we topically infect with HIV for up to 2h. However, these explants are prone to significant amounts of autofluorescence which impedes accurate analysis.

To assess the frequency and number of HIV+ DCs in an image we first ran AFid to generate a mask of autofluorescence within the image. We then used a spot counting algorithm^18^ to segment individual HIV particles, and subsequently segmented individual cells using their nuclei to generate single cell data. The fluorescent channels and masks of both the autofluorescence and HIV particles were then converted to FCS format for single cell analysis in FlowJo. Gating on putative CD11c+ DCs (**Fig. 3a, top left**) we then excluded autofluorescent cells, measured as the percentage of the cell body overlapping with the autofluorescence mask (**Fig.3a, top right**). After excluding cells containing autofluorescence we found that just 2% of DCs were HIV+ (contained at least 1 HIV particle) (**Fig. 3a., bottom right**) vs. 16% without excluding autofluorescent cells (**Fig 3a., bottom left**). This corresponded to a large difference in the total number of HIV+ DCs identified, 10 vs. 168 cells (**Fig. 3c**). We visually inspected the co-ordinates of putative HIV+ DCs after autofluorescence exclusion and were able to confirm that these cells corresponded to legitimate HIV-DC interactions as shown in the zoomed in images in **Fig. 3f**. In contrast, false positive signals were almost all due to autofluorescent signals spanning multiple channels. Importantly, we found that it was not possible to simply circumvent autofluorescence by gating specifically on true HIV+ DCs. This was because they did not occupy a unique position on biaxial plots and therefore could not be gated on without also including autofluorescent cells (**Fig. 3b**). Therefore highlighting the necessity of a computational approach to tag false positive signals within a mixed population.

**Fig. 3:**
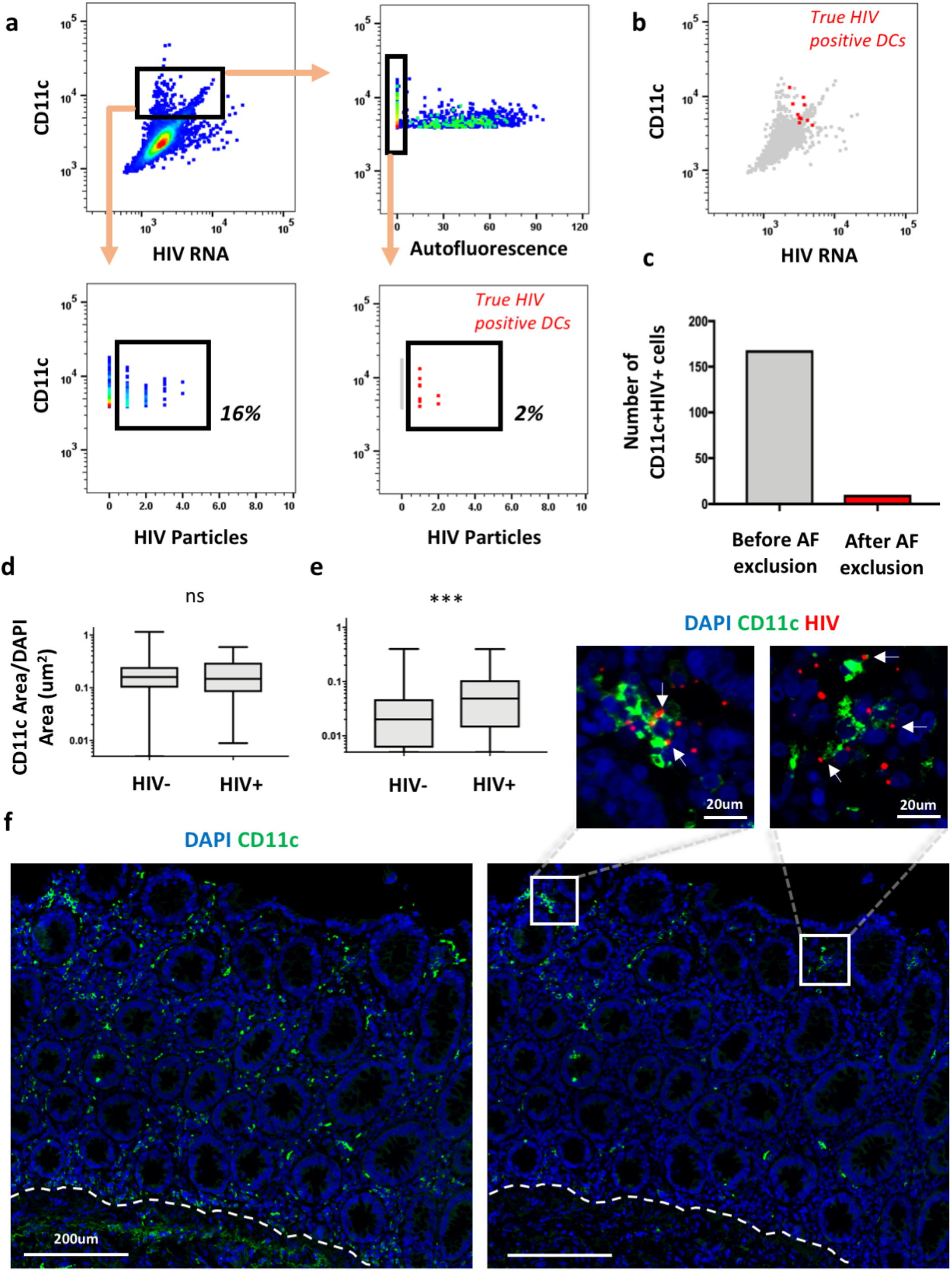
AFid facilitates analysis of early HIV-target cell interaction. Human colorectal explants were topically infected with HIVBal for 30min, fixed, sectioned and then stained for CD11c, HIV RNA and DAPI. **a**, Single cell segmentation and HIV spot segmentation were performed followed by FCS file generation for analysis using FlowJo. The figure shows a gating strategy for identifying CD11c+ cells that contain at least one HIV particle, either with, or without first gating-out cells containing autofluorescence. The autofluorescence parameter shows the percentage of the cell body overlapping with the autofluorescence mask generated by AFid. **b**, Overlay of CD11c+HIV+ cells after gating out autofluorescence, showing that they cannot be specifically gated-on without first excluding autofluorescent cells. **c**, Total number of CD11c+HIV+ cells before and after autofluorescence exclusion, as gated on in part a. **d,e**, A whole tissue image from one donor was divided into 100×100um^2^ quadrats, each classified as HIV- or HIV+, and CD11c labelling was measured before (d) and after (e) autofluorescence exclusion. CD11c expression was measured per um^2^ of DAPI. Quadrats with DAPI staining less than 1/10^th^ their area (non-tissue areas) were excluded. Boxplots show the min, first quartile, median, third quartile and max values. **f,** A cropped area of a whole-tissue image of HIV-infected colorectum before (left) and after (right) autofluorescence exclusion from the CD11c channel. Zoomed images of the boxes show interactions of HIV with CD11c+ cells (white arrows) in the image after autofluorescence exclusion. The broken white line indicates the base of the lamina propria. A two-tailed Mann Whitney test was performed in part **a** and **b**. ns = not significant; ***p=0.0002.

Finally, we show that the presence of autofluorescence can mask spatial relationships which are then revealed by its exclusion. DCs are highly migratory and thought to be an important early target cell for HIV transmission^19^. To assess whether DCs may be specifically localising to HIV early post-exposure we divided an image into 100×100um^2^ quadrats and classified each as HIV− or HIV+ and then measured the density of CD11c labelling in each area to quantify expression. Prior to autofluorescence exclusion (setting pixel values to 0), the apparent CD11c expression did not significantly differ between HIV− and HIV+ areas (**Fig. 3d**), whereas after exclusion CD11c expression was revealed as significantly higher in HIV+ areas compared to HIV-areas (**Fig. 3e**). This was due to a large amount of measured CD11c expression being derived from autofluorescence (**Fig. 3f)**. Further, we found that CD11c, HIV and autofluorescent cells were differentially located. CD11c and HIV clustered toward the tip of the lamina propria where the majority of interactions took place (**Fig. 3f, zoomed images**), whilst autofluorescent cells were particularly clustered toward the base of the lamina propria, thus skewing the results. Together these data demonstrate how the presence of autofluorescence can drastically skew measurements, and that its post-acquisition exclusion, here using AFid, can enable accurate image analysis.

## Discussion

Here we have presented AFid, a first of its kind method for identifying autofluorescence in multi-channel fluorescence microscopy images post-acquisition. We have shown that AFid is able to identify autofluorescence across tissue types, staining panels and image acquisition conditions. Importantly, we showed that the presence of autofluorescence can have a major impact on downstream analyses which can be mitigated through the use of AFid.

Our rationale for creating AFid lay in its necessity for accurately measuring HIV-cell interactions in our images of HIV-infected FFPE human colorectal tissues. In particular, our staining protocols were incompatible with chemical bleaching methods discussed in the introduction of this paper^9,12–14^, and we did not have the necessary resources to perform spectral unmixing^15^ or background subtraction^2,16,17^. Noting that the majority of interfering autofluorescence in our samples were bright, spatially distinct objects appearing across multiple channels we reasoned that they could be identified and excluded from downstream analyses by way of a computational approach. Indeed, as demonstrated here the application of AFid was sufficient to enable accurate quantitative measurements of HIV-cell localisation.

AFid has several distinct advantages. First, it is a post-acquisition algorithm. This is important because the user does not need to pre-emptively deal with autofluorescence. Additionally, it can be applied to existing microscopy datasets potentially mitigating the need for optimisation and repeat experimentation. Indeed, this is exemplified in our own case, where bulk image data sets from HIV-infected explants were probed using RNAscope and multiple immune markers (partially presented in **Fig. 3**, unpublished data) but quantification hindered by autofluorescence. In this case, even if further lab-based optimisations proved successful at mitigating the impact of autofluorescence for our specific tissues, repeating the experiments would require the use of precious samples and a significant investment of time and resources.

The second key advantage of our algorithm is that its design allows for the generalised detection of autofluorescence across varying tissues and image acquisition conditions, as demonstrated here. In particular, we assume that due to the long excitation/emission wavelengths of autofluorescence^10^ (**Supplementary Fig. 1**), textural features of autofluorescent objects will exhibit a conserved topology across channels, and therefore occupy a unique position in the feature space, relative to other objects (as shown in **Supplementary Fig. 6–8**). Accordingly, the employment of clustering, here using *k* means with automated choice of *k*, is able to consistently isolate a cluster containing mainly autofluorescent objects, despite variations across tissues and image acquisition conditions.

Despite the advantages that have been discussed there are several limitations to our approach. The major limitation is that AFid cannot identify autofluorescence that is largely overlapping with real signal. This situation may occur if the object of interest is inherently autofluorescent or localised to a highly autofluorescent area of the tissue. This maybe an advantage or a disadvantage depending on the nature of the measurements to be performed. For example, AFid would aid in object counting measurements by retaining objects with mixed real-signal and autofluorescence, whilst excluding objects that purely contain autofluorescence (e.g. autofluorescent macrophages, **Fig. 2**). However, it would not be suitable if precise fluorescence intensity measurements were required of a stained object that contained some autofluorescence. Another potential limitation is that the algorithm requires that autofluorescent signals are present, even if only faintly, across at least two acquired channels. In our experience this was true of all bright interfering autofluorescence in the various tissues used for this study and fits in with the well-known broad spectra of autofluorescence^10^ (**Supplementary Fig. 1**). However we cannot discount the possibility of specific types of autofluorescence having narrow spectral profiles that might appear in only one channel. For the limiting cases discussed here, optimisation of chemical quenching^9,12–14^ or algorithmic background subtraction methods^2,15–17^ may be necessary.

Nonetheless, here it is demonstrated that AFid has major utility for quantitative measurements in human gut tissue stained for common immune markers, and have shown that it is able to identify autofluorescence in various other tissue types. With the rise of quantitative image analysis, particularly single cell cytometry pipelines as shown here, there is an increasing need for image processing algorithms to filter out artefacts and enable accurate measurements. Accordingly, AFid provides a major leap forward in the extraction of useful data from images plagued by autofluorescence by offering an approach that is easily incorporated into existing workflows in ImageJ, Matlab and R, and that can generalise to various samples, staining panels and image acquisition methods.

## Methods

### Immunofluorescence staining

Tissues were fixed in 4% paraformaldehyde (Electron Microscopy Sciences) for 18-24h at room temperature then immersed in 70% ethanol prior to paraffin embedding. 4um paraffin sections were adhered to glass slides (SuperFrost Plus, Menzel Glazer), baked at 60°C for 40 min, dewaxed in xylene followed by 100% ethanol then air dried. All wash steps described herein were carried out by immersing slides in three successive Coplan Jars of Tris-buffered saline (Amresco, Cat: 0788) on a rotator for a total of 10 minutes. Antigen retrieval was then performed using a pH9 antigen retrieval buffer (DAKO) in a decloaking chamber (Biocare) for 20 min at 95°C. Slides were then washed in TBS. To acquire unlabelled background images (**Fig. 2, Supplementary Figs. 2 and 9**), sections were stained with 1ug/ml DAPI (Roche) for 3 minutes, mounted under coverslips with SlowFade-Diamond Antifade (Molecular Probes) and the whole section imaged on an Olympus VS120 microscope (see Image acquisition below). Coverslips were then floated away in TBS and sections on slides were blocked for 30 min (0.1% saponin, 1% BSA, 10% donkey serum, diluted in TBS) at room temperature. Sections were then washed in TBS and incubated with primary antibodies overnight at 4°C. Antibodies for primary detection include: Abcam: - rabbit CD11c (EP1347Y), mouse CD3 (F7.2.38), rabbit CD8 (polyclonal, ab4055); DAKO – rabbit CD3 (polyclonal, A045229-2); Affinity Biologicals – sheep FXIIIA (polyclonal). Sections were then washed in TBS and incubated with secondary antibodies for 30min at room temperature. Donkey secondary antibodies (Molecular Probes) against rabbit, mouse or sheep were used and were conjugated to either Alexa Fluor 488 or 546. Sections were stained with DAPI (if not already performed in a previous step) and mounted with SlowFade-Diamond Antifade.

### HIV explant infection

Healthy Inner foreskin explants were infected with either HIV_Bal_ or Transmitted/Founder HIV-1 Z3678M using an explant setup as previously described^20^. A TCID_50_ of 3500 (titrated on TZMBLs as previously described^21^) was used to infect all explants. Tissues were then fixed and paraffin embedded as described above.

### RNAScope

Detection of HIV RNA was performed using the ‘RNAscope 2.5HD Reagent Kit-RED’ and following the manufacturer’s protocol (Cat: 322360, ACD Bio) with custom probes (consisting of 85 zz pairs) against HIV-1_BaL_ (REF: 486631, ACD Bio) spanning base pairs 1144-8431 of HIV-1_BaL_ sequence. Following the RNAscope protocol, sections were stained from the blocking step as detailed above.

### Microscopy

Imaging was performed using an Olympus VS120 Slide Scanner with ORCA-FLASH 4.0 VS: Scientific CMOS camera. VS-ASW 2.9 Olympus software was used for acquisition of images and conversion of raw vsi files to tiff format for downstream processing. Objectives used are indicated in figure legends and include: ×10 (UPLSAPO 10×/ NA 0.4, WD 3.1 / CG Thickness 0.17), ×20 (UPLSAPO 20×/ NA 0.75, WD 0.6 / CG Thickness 0.17) and ×40 (UPLSAPO 40×/ NA 0.95, WD 0.18 / CG Thickness 0.11–0.23). Channels used include: DAPI (Ex 387/11-25 nm; Em: 440/40-25 nm), FITC (Ex:485/20-25 nm; Em: 525/30-25 nm), TRITC (Ex:560/25-25 nm; Em: 607/36-25 nm) and Cy5 (Ex: 650/13-25 nm; Em: 700/75-75 nm). For ×40 images, Z-stacks were acquired 3.5um above and below the plane of focus with 0.5um step sizes. Huygens Professional 18.10 (Scientific Volume Imaging, The Netherlands, http://svi.nl) CMLE algorithm, with SNR:20 and 40 iterations, was used for deconvolution of Z-stacks. For images where the unstained background was acquired prior to staining, images were aligned using the ImageJ plugin multiStackReg vs1.45 with the DAPI channel serving as a reference for alignment.

### Acquisition of Autofluorescence Spectra

Autofluorescence spectra of unstained tissue samples (**Supplementary Fig. 1**) were acquired using an Olympus FV1000 laser scanning confocal microscope with a ×20 objective. The excitation lasers lines 405nm, 473nm and 559nm were used and emission spectra were acquired using a 20nm wide bandpass filter, shifted in 20nm intervals from 415-795nm, 490-790nm and 575-795nm respectively.

### Generation of Intersection Mask

A mask of the intersection of the two channels was used for autofluorescence removal. This is termed the ‘intersection mask’. The intersection mask contains only signals present in both channels and therefore contains the autofluorescent ROIs among other objects such as co-stained markers and dim background stromal fluorescence. The intersection mask was generated by the following procedure. Each channel was Gaussian blurred with a sigma of 2. A Niblack threshold was then applied to each channel (threshold radius 30 pixels) to generate binary masks. The intersection (‘AND’ operation) of these masks was then taken and used for autofluorescence classification by clustering as detailed below.

### Clustering for autofluorescence identification

Within the objects defined by the intersection mask we measured multiple features in each of the two channels on non-Gaussian blurred images. These features included standard deviation, kurtosis, as well as the inter-channel Pearson’s correlation coefficient of corresponding pixels. These features were transformed by taking the natural log (standard deviation, skewness and kurtosis) or the inverse tanh transformation (correlation). All features were standardised by dividing by the standard deviation of the transformed feature values. *k*-means clustering was then performed on these features to identify a cluster of ROIs which are likely to be autofluorescent. The cluster with the highest average correlation value was defined as the cluster containing autofluorescent ROIs. A well-chosen number of clusters (*k*) is important for detecting a homogeneous cluster of autofluorescent ROIs. As such we developed an automated approach for optimal choice of *k* (high sensitivity and specificity). The procedure is as follows. 1. *k*-means is performed iteratively with 3-20 clusters 2. A two-tailed t-test is performed on the arctanh transformed correlation values of the two clusters with highest average correlation values. 3. The test statistic values are then plotted against *k*, which produces an asymptotically decreasing function (**Supplementary Fig. 13**). 4. We developed an ‘elbow method’ approach to finding the optimal cluster number. A straight line is drawn connecting the statistic value for the lowest *k*, to that of the highest *k*. The perpendicular distance of each plotted point to the line is measured and the optimal *k* is estimated to correspond to the point with the greatest distance below the line. This method is illustrated in (**Supplementary Fig. 13**). The intersection mask is then modified, keeping only the objects identified as autofluorescence.

### Custom dilation function to outline autofluorescent ROIs

After clustering and creating a mask of autofluorescent objects we then employed a custom dilation function to outline the full body of autofluorescent objects for removal. The essence of the algorithm is to evenly distribute points within an amorphous object and then to expand out from these points in all directions until a halting condition is met.

To distribute points the following approach was developed: 1. ROIs in the autofluorescence mask were skeletonised, reducing objects to a line of 1 pixel-width that follows the morphological gradient of the original object. 2. End-node pixels for each object in the image were first identified, defined as having only one neighbour. If there were no end-nodes for an object, as in the case of an annulus, the top-left-most pixel was defined as the end-node. 3. A skeleton tracing algorithm was employed that starts from the end nodes and moves throughout the skeleton, distributing centres for expansion every 20 pixels (illustrated in **Supplementary Fig. 4d**). Tracing of pixels to neighbours occurred as long as the neighbouring pixel was in the skeleton and had not yet been traced by another point. Once these conditions were no longer met, tracing for a given object was halted. Expansion from distributed centres was carried out as follows: 1. Lines of length 60 pixels emanating from centres were drawn in all directions separated by an angle of theta where theta was defined by the law of cosines. 2. Pixel values of the Gaussian blurred image for each channel were measured beginning from the point of intersection of the line and perimeter of the object in the intersection mask, to the end of the line. 3. The co-ordinates of the first point where pixel values increased were recorded for each line. 4. A new outline of the object was created by combining these co-ordinates (**Supplementary Fig. 4e**). 5. Pixel values of the new outline of the object were set to 0.

### Assessment of custom dilation function robustness with varying parameters

This section specifically details the methods for generating data for **Supplementary Fig. 4g**, which assesses both the utility of the custom dilation function for capturing autofluorescent pixels and its robustness against varying parameters. For this figure whole-slide scanned images of colorectal tissue before antibody labelling and after CD11c/CD3 antibody labelling were used (data from **Fig. 2a-c).** AFid was run with or without use of the custom dilation function and using the CD11c and CD3 channels as inputs. This was performed for varying threshold radii and for varying cluster number *k* as shown in the graph. To assess coverage of autofluorescent pixels (true positive) and real signal pixels (false positive) upon parameter variation as shown in the graph, we generated ground-truth masks. To do this, we specifically used the CD3 channel due to the consistently high density of T cells across images, thus providing enough data to assess the false positive rate. We generated a mask of the unlabelled background image (autofluorescence only) and the CD3 channel after staining (CD3+ T cells and autofluorescence). Our aim was to use the background image to parse out the autofluorescent pixels contained within the CD3 channel image.

First, we performed a morphological watershed on the CD3 channel mask to separate touching T cells and autofluorescence. We then performed a binary reconstruct of the watershed CD3 mask with the background image mask. This generated an estimate mask of the autofluorescent pixels within the CD3 image which we used as a ground-truth for autofluorescence. We then generated the real-signal ground-truth mask by subtracting the autofluorescence ground-truth mask from the CD3 channel mask. The ground-truth masks were then manually inspected and objects were removed which did not clearly correspond to T cells in the real-signal mask or autofluorescence in the autofluorescence mask.

### Algorithm performance assessment

The performance of our algorithm was tested using three different staining panels on human colonic tissue as shown in **Fig. 2**. To benchmark performance assessment, we manually annotated regions of the intersection mask (see ‘Generation of Intersection Mask’ above) as belonging to ‘real’ or ‘autofluorescent’ signals. Delineation of the two types of signal was achieved using the ‘unstained background image’ as a reference (see ‘Immunofluorescent Staining’ above). In total 400 ROIs, 200 for each category, were annotated. The actual annotation was performed using the Cell Counter Plugin in ImageJ. Results were exported as a csv file, where each row indicated an individual ROI, its category and x,y co-ordinates.

The two fluorescent channels, intersection mask and spreadsheet of annotated ROI co-ordinates were fed in to R. *k*-means clustering with estimated *k* was then performed as described above. The true positive rate and false positive rate were thus determined as the proportion of ROIs in each category that resided in the ‘autofluorescence cluster’, which was the cluster with highest average correlation values (**Supplementary Fig. 5 and 13**).

### HIV spot segmentation

Spot counting was performed using a custom MATLAB script implementing the spot counting technique presented by Battich et al. (2013). First, a manual threshold was performed on the HIV RNA channel to identify areas that have HIV stain present. The *IdentifySpots2D* function by Battich et al. was then used to identify the number of spots, with the detection threshold set to a generous value of 0.01 and the number of deblending steps equal to 2. Finally, any spots identified were excluded if they were not present in the threshold mask obtained previously.

### Single cell segmentation

To perform single cell segmentation, a custom MATLAB script was used. Briefly, a Gaussian filter with a full-width at half maximum of 10 pixels was applied to the DAPI image to ensure that each nucleus has only one locally maximum pixel intensity.

Further, the *imordfilt2* function is used to ensure that maxima are not less than 7 pixels apart. Watershed segmentation is performed using the *watershed* function to identify nuclear boundaries. Objects with diameters less than 10 pixels or greater than 50 pixels were discarded, and the nuclear objects are dilated by 6 pixels to estimate the cell body. The *regionprops* function was finally used to measure the mean pixel intensities of other image channels within each identified cell boundary, as well as the number of HIV RNA spots identified within each cell. The data was exported as a *.csv* and was analysed using FlowJo.

## Code availability

This algorithm has been implemented with user interfaces in Fiji (Madison Version), R (3.5.3) and Matlab (r2017b) to accommodate the diverse image analysis community. The code and user documentation are available **https://ellispatrick.github.io/AFid.**

## Data availability

Data sets used in this study are available from the corresponding author upon reasonable request.

## Ethics for use of human tissue samples

This study was approved by the Western Sydney Local Area Health District (WSLHD) Human Research Ethics Committee (HREC); reference number (4192) AU RED HREC /15 WMEAD/11. Human colorectal and skin tissues used for this study were approved by this committee and all patients were consented prior to sample collection. Brain and Heart tissue images were donated data.

## Acknowledgements

We would like to acknowledge the Cell Imaging Core Facility at the Westmead Institute for their support and services. Also, Rashid F., member of the Chong Lab, for the rat heart images used for this study, and Bright F., for the human brain tissue images used for this study. Also a special thanks to Cantrill L. for reviewing this manuscript. This work was funded by the National Health and Medical Research Council (Australia).

## Author contributions

H.B conceived of the project, collected data and wrote the first implementation of the algorithm. N.C wrote the majority of the codebase in ImageJ, MATLAB, and R. K.M.B, K.J.S, T.L.C., A.N.H and E.P were intellectual contributors to the project. All authors were involved in writing and editing the manuscript.

---

**Supplementary Figure 1:**
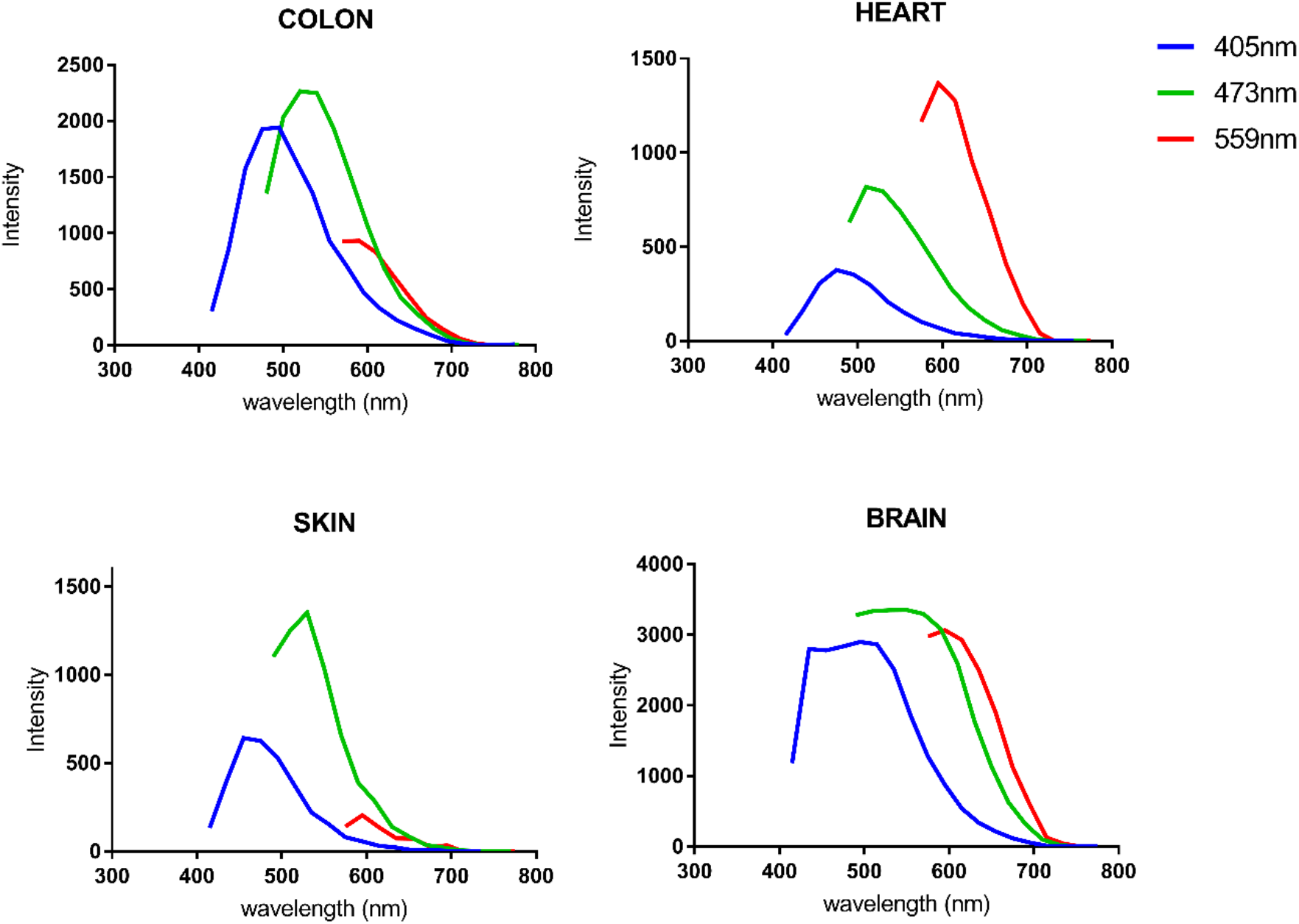
Excitation and emission spectra of autofluorescence in various tissues. The intensity of pixels corresponding to autofluorescent structures measured at 20nm intervals upon excitation with laser lines 405nm, 473nm or 559nm in human colon, skin and brain tissues, as well as rat heart tissue. Results shown as the intensity of the autofluorescent object minus the intensity of the stromal background for each wavelength. Results are shown for a single image for each tissue type.

**Supplementary Figure 2:**
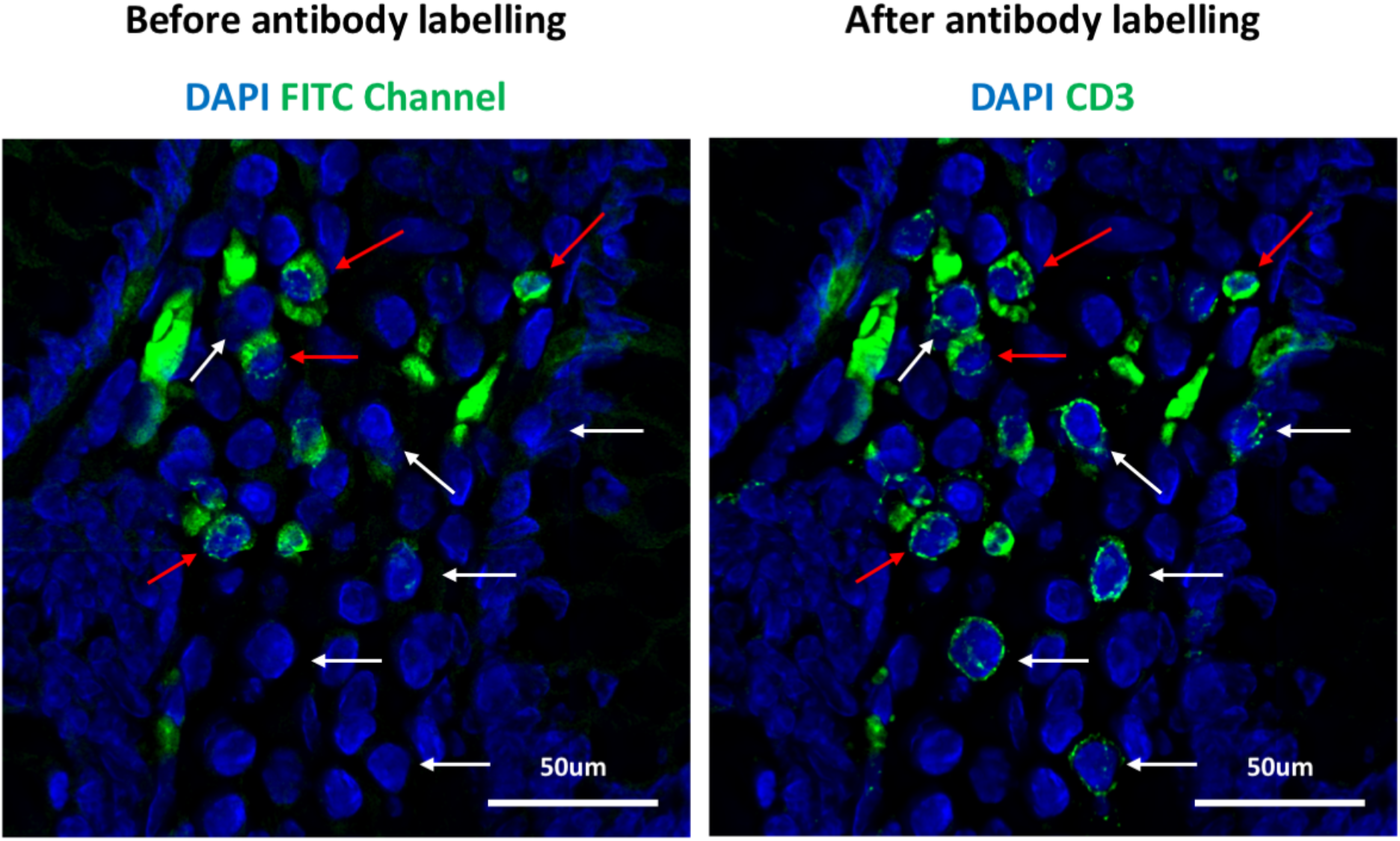
Autofluorescence inhibits assessment of CD3 labelling in the human colorectum. Fixed human colorectal tissue sections imaged prior to (left) and after labelling with mouse anti human CD3 and donkey anti-mouse AF488 (right). Red arrows indicate some autofluorescent cells and white arrows indicate CD3+ cells. Images are representative of 6 unique donors where CD3 staining was performed.

**Supplementary Figure 3:**
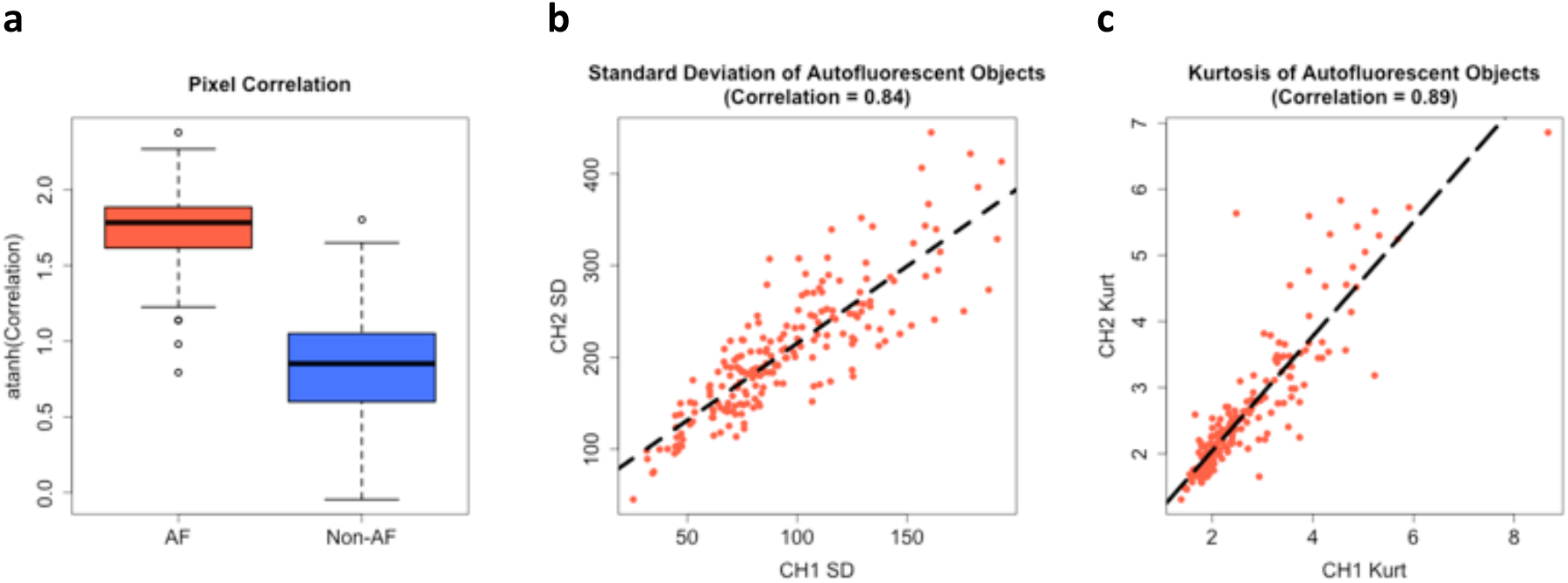
Features of autofluorescence are highly correlated between fluorescent channels. Fixed human colorectal tissue sections were stained for mouse anti CD3 and rabbit anti CD4, detected using donkey anti-mouse AF488 and donkey anti-rabbit AF546 respectively. An intersection mask was created using the two fluorescent markers (**Fig. 1**) and measurements performed on objects in the intersection mask. An unstained background image was used as a reference to manually annotate autofluorescent objects in the stained image. **a**, The arctanh transformed Pearson’s correlation coefficient values of autofluorescent objects vs non-autofluorescent objects within the intersection mask. The boxplots contain data from thousands of individual objects for each category. Boxplots show the min, first quartile, median, third quartile and max values. **b,c**, standard deviation (SD) and kurtosis (Kurt) measurements of autofluorescent objects in each channel (CH1 and CH2) used to create the intersection mask. A subsample of 200 autofluorescent objects, among thousands, is shown. A line of best fit is shown for both graphs. These graphs are representative of 13 total images used for this work.

**Supplementary Figure 4:**
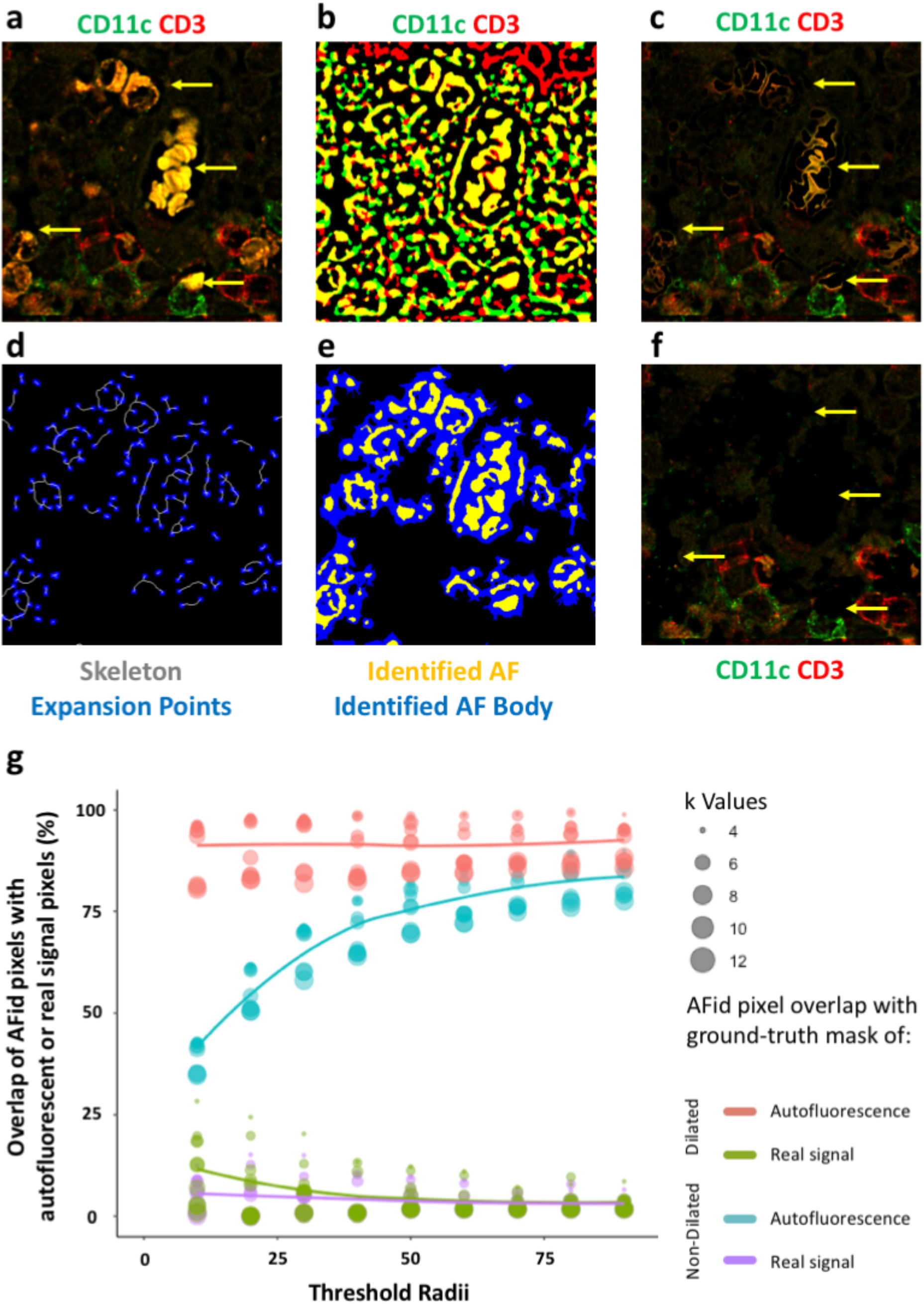
Custom dilation function to estimate the correct perimeter of autofluorescent ROIs. **a**, Fixed colorectal tissue sections were stained for rabbit CD11c and mouse CD3, followed by donkey anti rabbit AF488 and donkey anti mouse AF546. **b**, Fluorescent channels used to detect CD11c and CD3 were thresholded, binary masks created and the composite image displayed. Yellow indicates the overlapping area corresponding to the intersection mask. **c**, Image from part a with the pixels in the intersection mask from part b set to 0. **d**, Identified autofluorescent objects within the intersection mask in part b are skeletonised and points for outward expansion (blue) are distributed along the skeleton every 20 pixels. **e**, Thousands of equiangular lines are drawn outwards from the expansion centres identified in part d, each line propagating until it encounters a pixel brighter than the previous pixel, as measured in either the CD11c or CD3 channel. A mask of the identified autofluorescence body is thus generated for each fluorescent channel. **f**, Pixels corresponding to the identified AF body in each channel in part e, are set to 0. Yellow arrows indicate some autofluorescent objects. **g**, Measurement of overlap of autofluorescent pixels defined by AFid, with ground-truth masks of autofluorescence and real signal (CD3 antibody labelled) pixels. Measurements were performed with (red and green lines) and without (blue and purple) employing the custom dilation function and for varying cluster number *k* and varying threshold radii. Representative data of 3 images is shown. See methods for additional details.

**Supplementary Figure 5:**
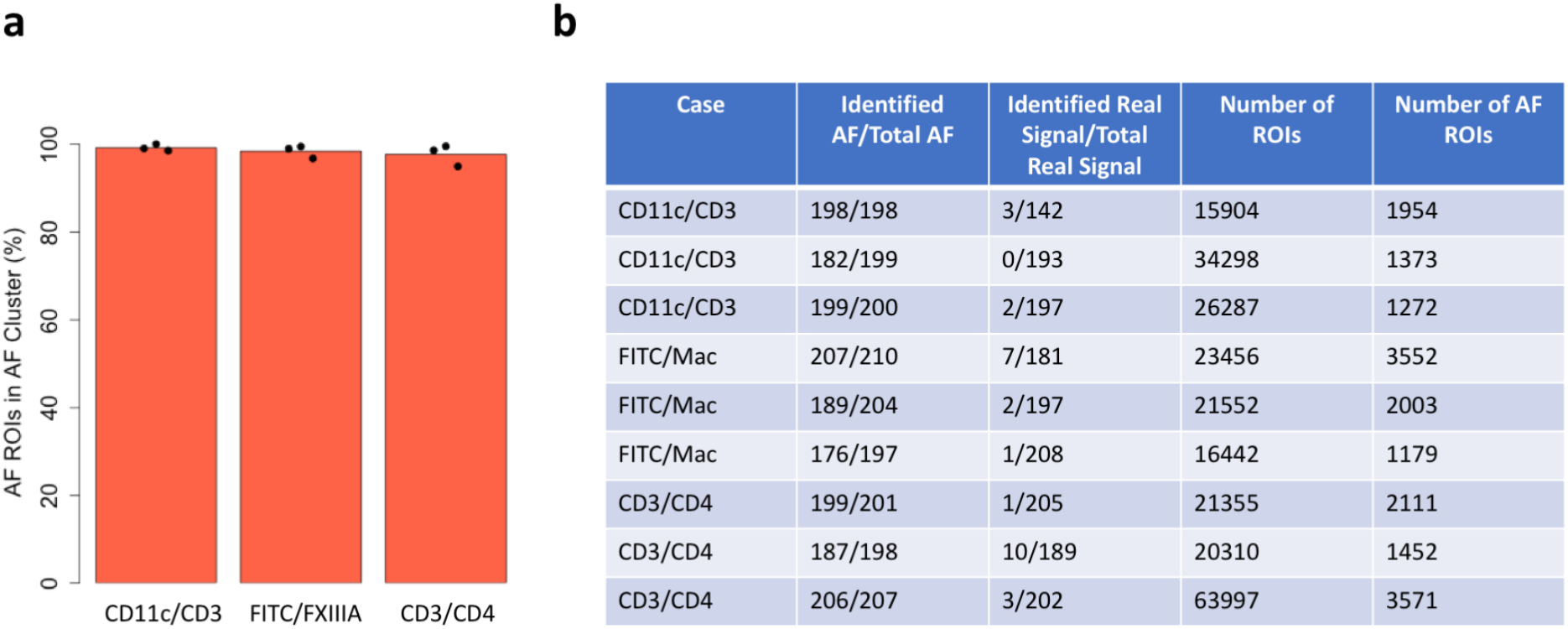
Specificity and sensitivity of autofluorescence removal for various use-cases in Figure 2. Fixed human colorectal sections were imaged prior to, and after labelling with antibodies against markers for three separate panels, CD11c/CD3, FITC/FXIIIA and CD3/CD4. FITC indicates an unstained open channel that was imaged. An intersection mask was created using the two fluorescent channels for each panel (as in Figure 1). Textural features of objects within the intersection mask were then measured for each channel, including standard deviation, kurtosis, as well as the inter-channel Pearson’s correlation coefficient of corresponding pixels. *k*-means clustering was then performed using these features and the cluster with the highest average correlation values was defined as the cluster containing autofluorescent ROIs. A ground truth for the classification of objects as autofluorescence or real signals (stemming from antibodies) was established by manually annotating a subset of up to 200 ROIs each, using the unlabelled background image as a reference. **a**, percentage of the ‘autofluorescence cluster’ comprised of autofluorescent ROIs (specificity), where the total number of ROIs in the cluster is defined as the sum of autofluorescent ROIs and ROIs stemming from real signal. Each data point represents counts performed on a unique donor for each panel. Mean values across the three donors are indicated above each column. **b**, table summarising the proportion of manually annotated autofluorescence or real signal assigned to the ‘autofluorescence cluster’ (sensitivity). The last two columns indicate the total number of ROIs and the total number of ROIs classified as autofluorescence respectively. Each row corresponds to results for unique donors for each use-case.

**Supplementary Figure 6:**
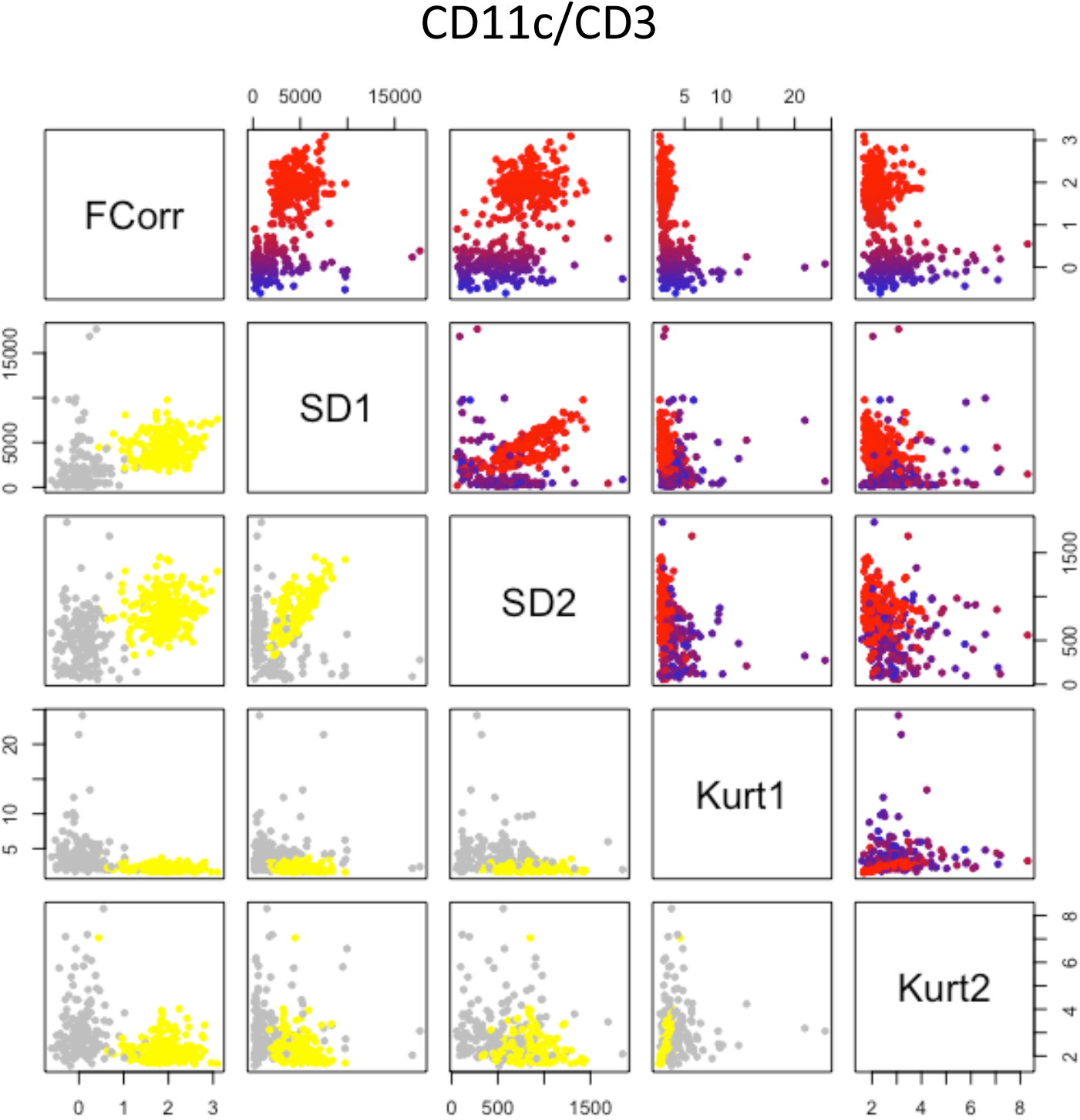
Pairwise plots of textural features used for *k*-means clustering of the non-co-expressed markers use-case. ROIs from *k*-means clustering on the CD11c/CD3 use-case in Supplementary Figure 5 are shown. In the bottom half, ROIs in the autofluorescence cluster are coloured yellow, whilst non-autofluorescent ROIs (real signal + dim stromal background fluorescence) are coloured grey. The top half shows paired plots as a heatmap of correlation values. FCorr = Arctanh transformed Pearson’s correlation coefficient values. SD1, SD2, Kurt1, Kurt2 = Standard deviation or Kurtosis values of ROIs in channels 1 or 2. The plot is representative of clustering performed on 3 independent donors.

**Supplementary Figure 7:**
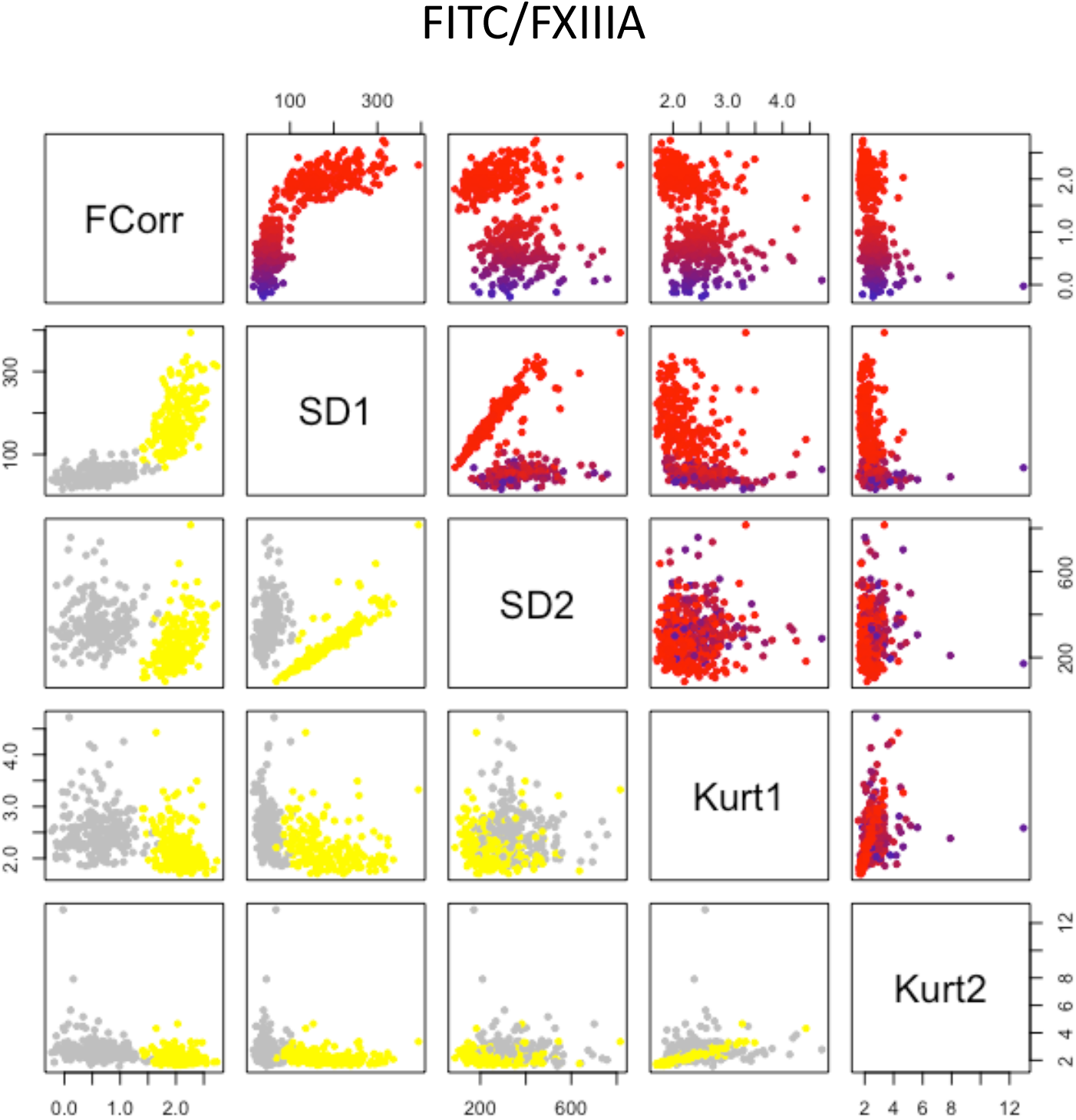
Pairwise plots of textural features used for *k*-means clustering of the autofluorescent cells use-case. ROIs from K-means clustering on the FITC/FXIIIA use-case in Supplementary Figure 5 are shown. In the bottom half, ROIs in the autofluorescence cluster are coloured yellow, whilst non-autofluorescent ROIs (real signal + dim stromal background fluorescence) are coloured grey. The top half shows paired plots as a heatmap of correlation values. FCorr = Arctanh transformed Pearson’s correlation coefficient values. SD1, SD2, Kurt1, Kurt2 = Standard deviation or Kurtosis values of ROIs in channels 1 or 2. The plot is representative of clustering performed on 3 independent donors.

**Supplementary Figure 8:**
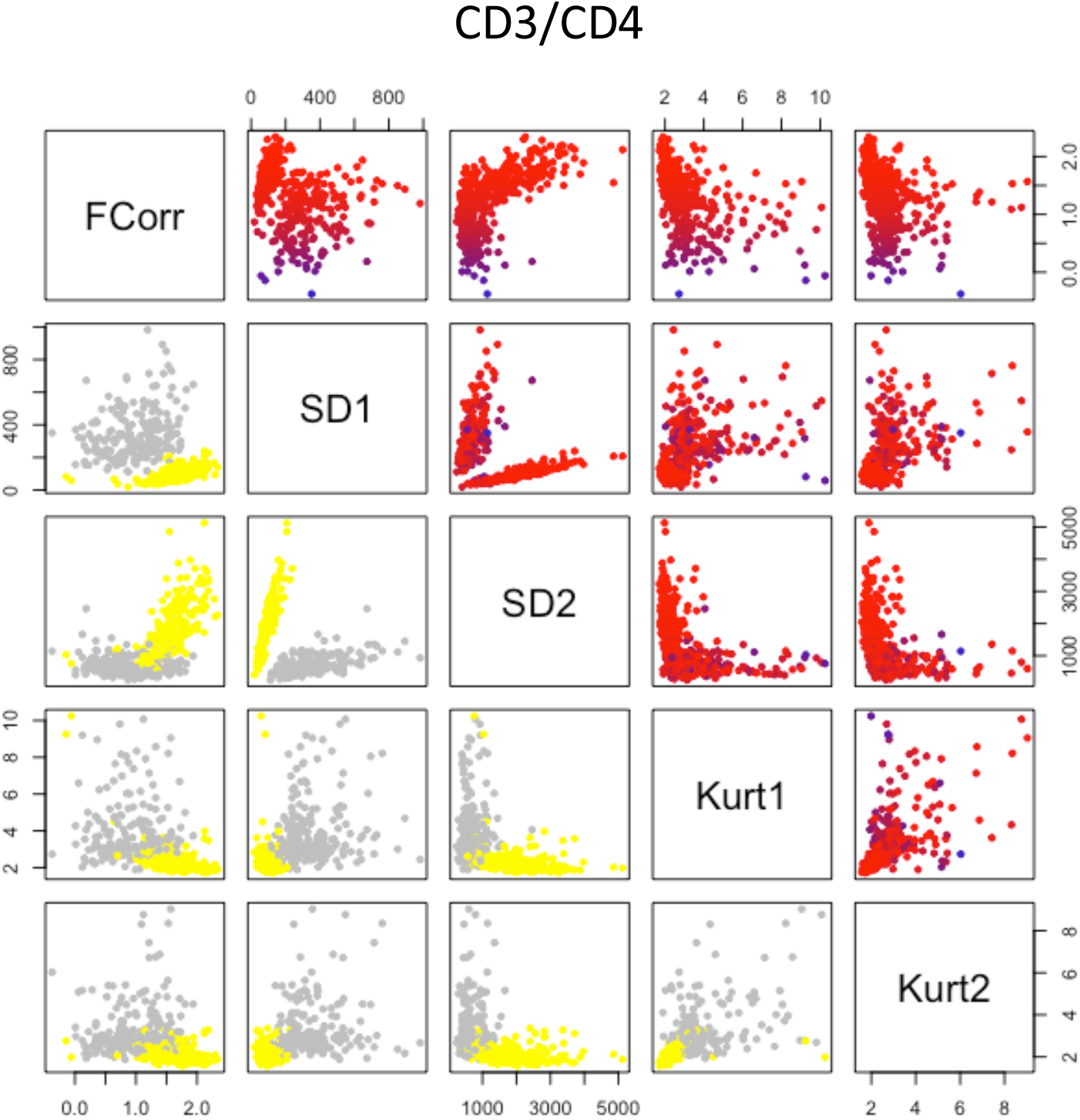
Pairwise plots of textural features used for *k*-means clustering of the co-expressed markers use-case. ROIs from K-means clustering on the CD3/CD4 use-case in Supplementary Figure 5 are shown. In the bottom half, ROIs in the autofluorescence cluster are coloured yellow, whilst non-autofluorescent ROIs (real signal + dim stromal background fluorescence) are coloured grey. The top half shows paired plots as a heatmap of correlation values. FCorr = Arctanh transformed Pearson’s correlation coefficient values. SD1, SD2, Kurt1, Kurt2 = Standard deviation or Kurtosis values of ROIs in channels 1 or 2. The plot is representative of clustering performed on 3 independent donors.

**Supplementary Figure 9:**
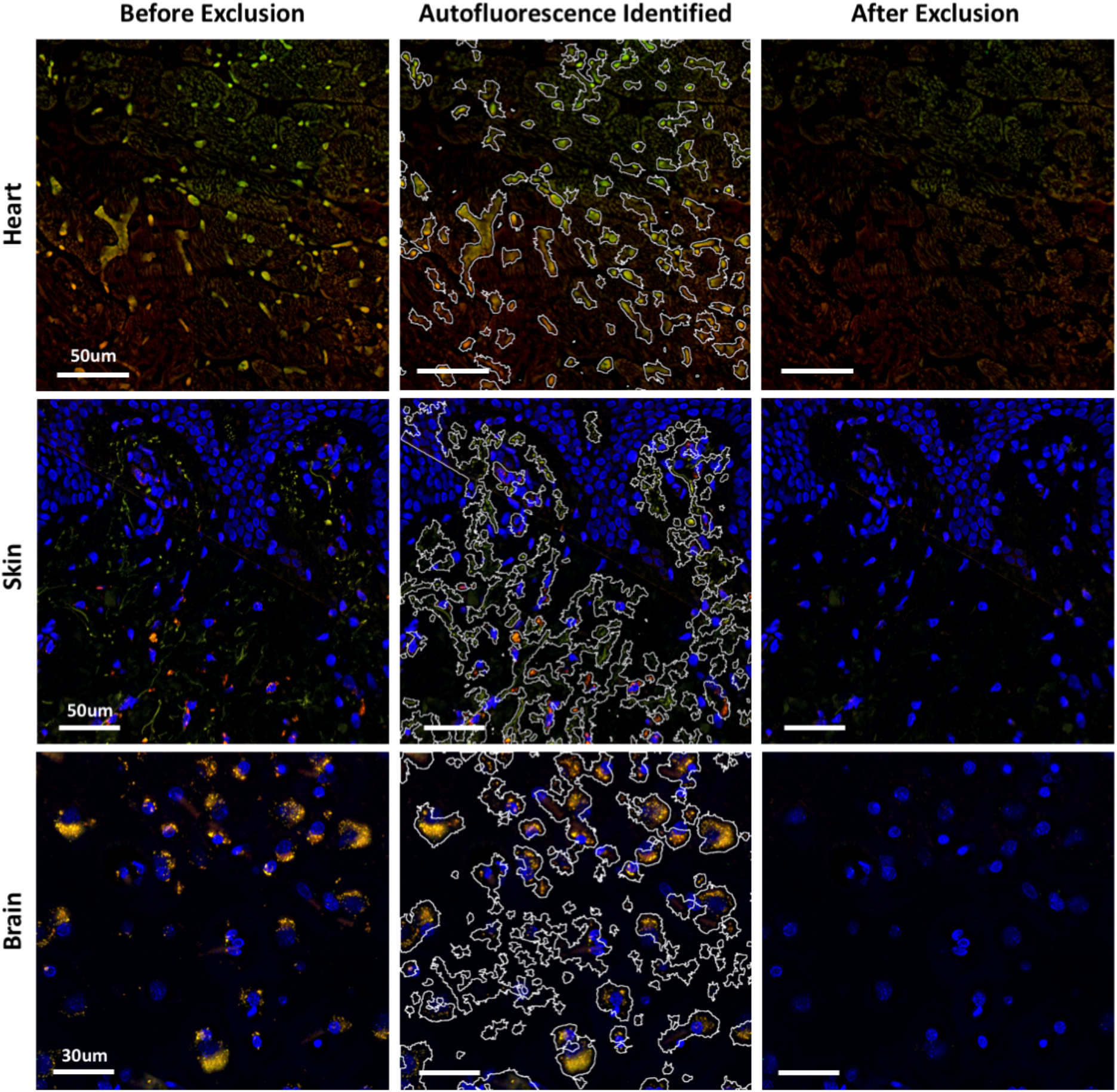
AFid performance across various tissue types. Unlabelled sections of fixed rat heart, human abdominal skin and human brain tissue are shown, prior to autofluorescence exclusion (left panel), with the boundary of autofluorescent structures identified by the algorithm (middle panel), and after exclusion by setting pixel values to 0 (right panel). Note that the images the skin and brain samples were counterstained with DAPI for visualisation. The images are representative areas from one whole-tissue image for each tissue type that was used for autofluorescence identification and exclusion.

**Supplementary Figure 10:**
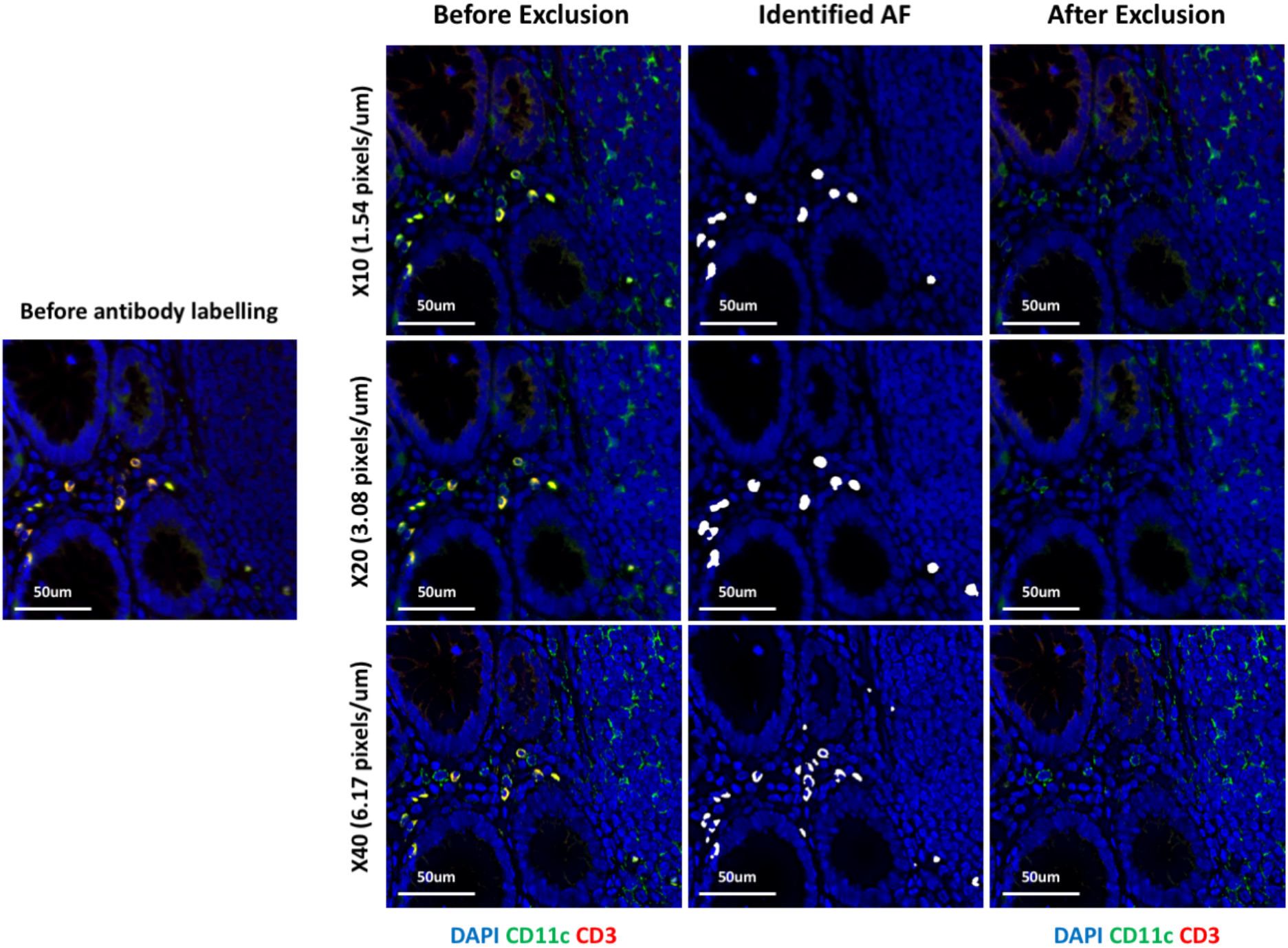
AFid performance at varying image resolutions. Fixed colorectal tissue sections were imaged prior to labelling (left, unlabelled image) and after labelling for rabbit anti CD11c and mouse anti CD3, followed by donkey anti rabbit AF488 and donkey anti mouse AF546. Images of the same area were taken with ×10, ×20 and ×40 objectives with an image resolution of 1.54, 3.08 and 6.17 pixels per um respectively. Images before autofluorescence removal (left panel), with a mask of the identified autofluorescence overlaid (middle panel) and after removal by setting pixel values to 0 (right panel) are shown. Images are representative of 3 unique donors where CD11c/CD3 staining was carried out and imaged at various magnifications.

**Supplementary Figure 11:**
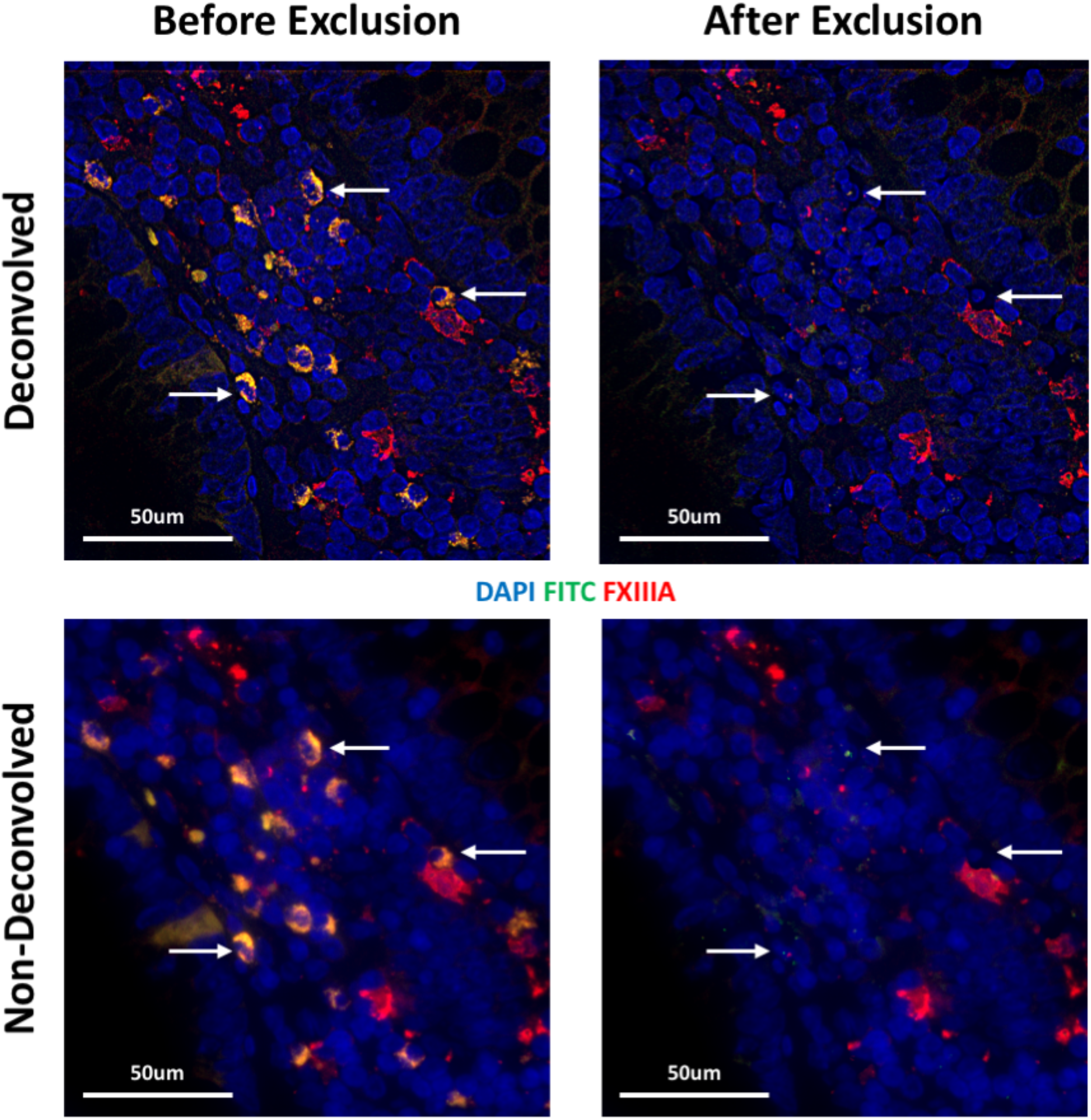
AFid performance before and after deconvolution. Fixed colorectal tissue sections were labelled with a sheep anti FXIIIA antibody followed by donkey anti sheep AF546. ‘FITC’ is the FITC channel, which was imaged but not used to detect any markers. Images are shown of the same area before (bottom row) and after (top row) deconvolution using Huygens deconvolution software, CMLE algorithm. Images before (left panel) and after (right panel) autofluorescence exclusion are shown. White arrows indicate some autofluorescence identified by the algorithm. Images are representative of 3 unique donors stained for FXIIIA and processed using AFid before and after deconvolution.

**Supplementary Figure 12:**
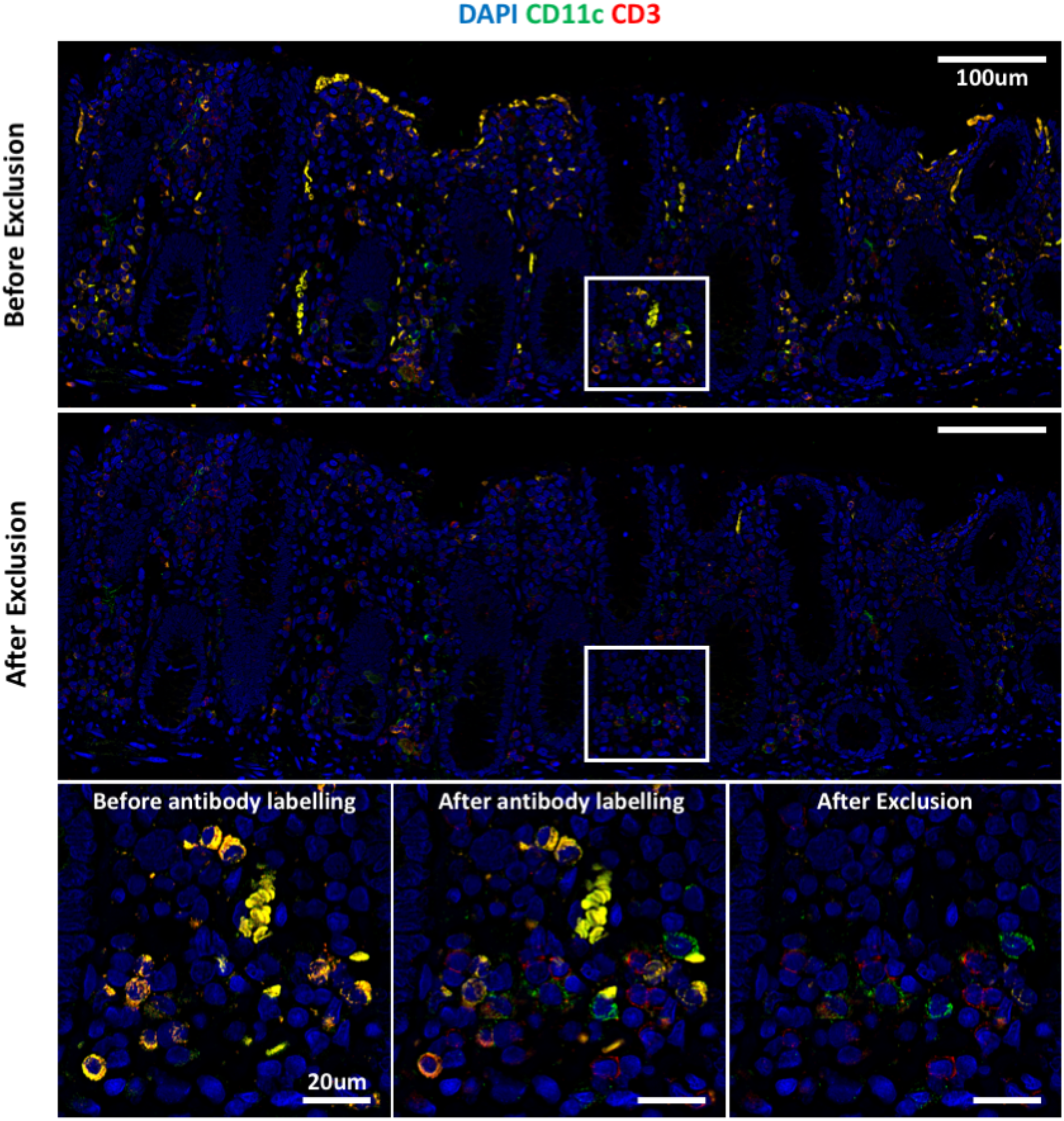
AFid performance on large images. Fixed colorectal tissue sections were imaged prior to and after labelling for rabbit anti CD11c and mouse anti CD3 antibodies, followed by donkey anti rabbit AF488 and donkey anti mouse AF546. A large area of tissue was imaged and the results before (top panel) and after (middle panel) autofluorescence exclusion are shown. Zoomed in images of the area outlined are shown with an additional image of the unlabelled section outlining the distribution of autofluorescence. Image is representative of 3 unique donors where CD11c/CD3 staining was carried out.

**Supplementary Figure 13:**
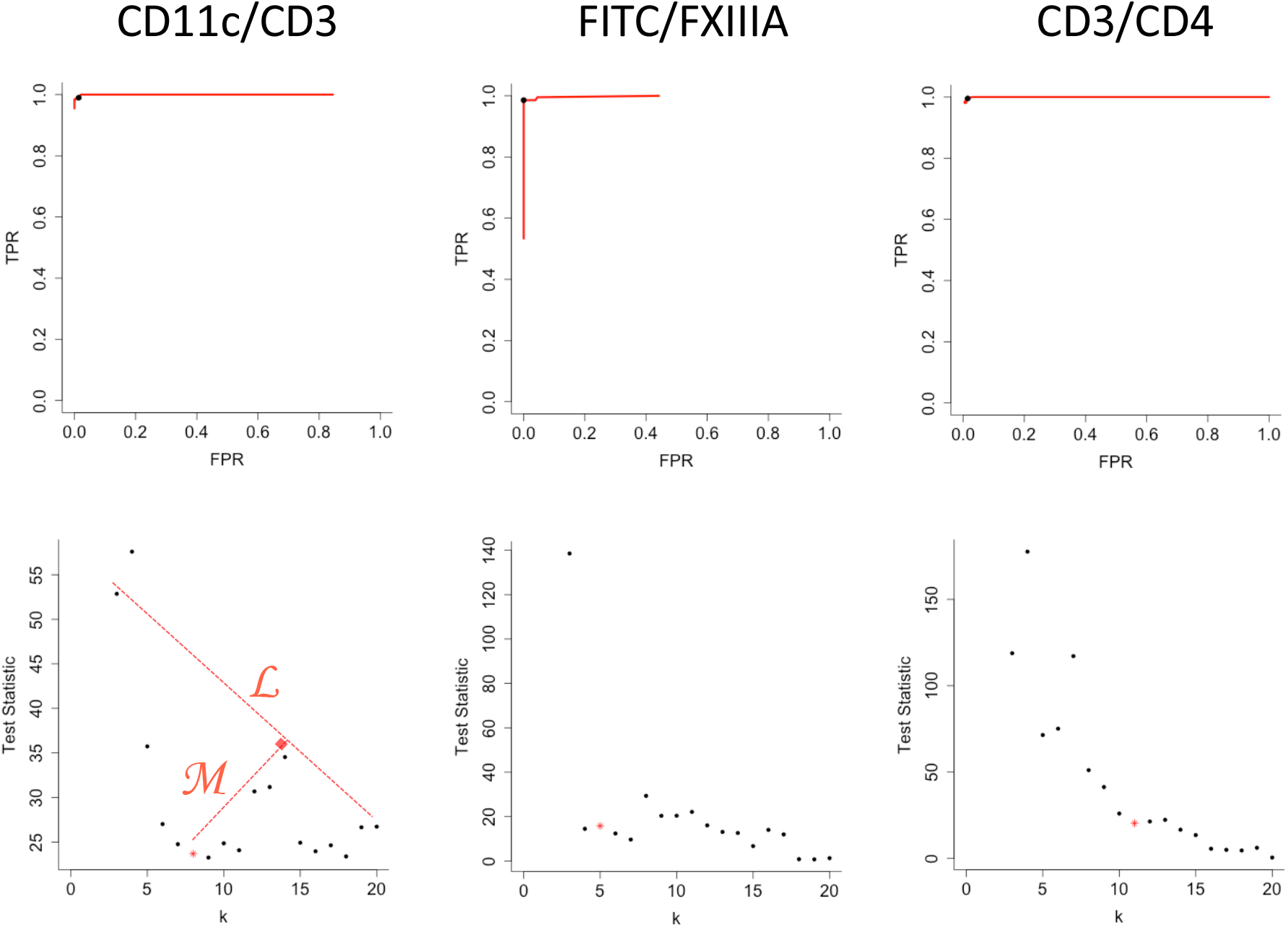
Estimation of optimal cluster number for *k*-means. *k*-means clustering as described in supplementary figure 5 was performed iteratively for 3-20 clusters and the distribution of paired true positive rate (TPR) and false positive rate (FPR) values for each cluster number is indicated by the red line for each use-case (top row). A high TPR or FPR corresponds to a high proportion of the ‘autofluorescence cluster’ comprising manually annotated autofluorescence or real signals respectively. The bottom row shows plots of the cluster number *(k)* versus, test statistic of a t-test (two-tailed) comparing Pearson’s correlation coefficient values of clusters with the highest (‘autofluorescence cluster’) and second highest average correlation values. An elbow method approach for estimation of optimal K is illustrated in the bottom left plot. A line (L) is drawn between the first and last plotted values. The line M indicates the plotted value that is below the line L, and has the greatest perpendicular distance to that line. The cluster number corresponding to this plotted point is estimated as the optimal cluster number. The points with optimal cluster number for each plot are indicated as a red *. The TPR/FPR of the optimal cluster number for each use-case is indicated by the black dot in the top row of plots. Data are representative of analysis performed on 3 unique donors for each use-case.

## References

1. Zrazhevskiy, P. & Gao, X. Nat Commun 4, 1619 (2013).

2. Gerdes, M.J. et al. Proceedings of the National Academy of Sciences 110, 11982–11987 (2013).

3. McKinley, E.T. et al. JCI Insight 2, (2017).

4. Lin, J.-R., Fallahi-Sichani, M. & Sorger, P.K. Nature Communications 6, 8390 (2015).

5. Sorger, P.K. et al. eLife 7, e31657 (2018).

6. Gut, G., Herrmann, M.D. & Pelkmans, L. Science 361, eaar7042 (2018).

7. Goltsev, Y. et al. Cell 174, 968–981.e15 (2018).

8. del Castillo, P., Molero, M., Ferrer, J. & Stockert, J. Histochemistry 85, 439–440 (1986).

9. Baschong, W., Suetterlin, R. & Laeng, H.R. Journal of Histochemistry & Cytochemistry 49, 1565–1571 (2001).

10. Croce, A.C. & Bottiroli, G. European Journal of Histochemistry 58, 2461 (2014).

11. Wizenty, J. et al. Journal of immunological methods 456, 28–37 (2018).

12. Billinton, N. & Knight, A.W. Analytical Biochemistry 291, 175–197 (2001).

13. Neumann, M. & Gabel, D. Journal of Histochemistry & Cytochemistry 50, 437–439 (2001).

14. Davis, S.A. et al. Journal of Histochemistry & Cytochemistry 62, 405–423 (2014).

15. Dickinson, M., Bearman, G., Tille, S., Lansford, R. & Fraser, S. BioTechniques 31, 1272–1278 (2001).

16. Woolfe, F., Gerdes, M., Bello, M., Tao, X. & Can, A. IEEE Transactions on Image Processing 20, 1085–1093 (2011).

17. Pang, Z. et al. Microscopy Research and Technique 76, 1007–1015 (2013).

18. Battich, N., Stoeger, T. & Pelkmans, L. Cell 163, 1596–1610 (2015).

19. Wu, L. & KewalRamani, V.N. Nature reviews. Immunology 6, 859–68 (2006).

20. Kim, M. et al. PLOS Pathogens 11, e1004812 (2015).

21. Harman, A.N. et al. Blood 118, 298–308 (2011).

